# Learning from Longitudinal Data in Electronic Health Record and Genetic Data to Improve Cardiovascular Event Prediction

**DOI:** 10.1101/366682

**Authors:** Juan Zhao, QiPing Feng, Patrick Wu, Roxana Lupu, Russell A. Wilke, Quinn S. Wells, Joshua C. Denny, Wei-Qi Wei

**Affiliations:** Department of Biomedical Informatics, Vanderbilt University Medical Center, Nashville, TN, USA; Division of Clinical Pharmacology, Vanderbilt University Medical Center, Nashville, TN, USA; Department of Medicine, University of South Dakota Sanford School of Medicine, Sioux Falls, SD, USA; Department of Medicine, Vanderbilt University Medical Center, Nashville, TN, USA

**Keywords:** cardiovascular disease prediction, machine learning, deep learning, genetics, electronic health records

## Abstract

**Background:** Current approaches to predicting Cardiovascular disease rely on conventional risk factors and cross-sectional data. In this study, we asked whether: i) machine learning and deep learning models with longitudinal EHR information can improve the prediction of 10-year CVD risk, and ii) incorporating genetic data can add values to predictability.

**Methods:** We conducted two experiments. In the first experiment, we modeled longitudinal EHR data with aggregated features and temporal features. We applied logistic regression (LR), random forests (RF) and gradient boosting trees (GBT) and Convolutional Neural Networks (CNN) and Recurrent Neural Networks, using Long Short-Term Memory (LSTM) units. In the second experiment, we proposed a late-fusion framework to incorporate genetic features.

**Results:** Our study cohort included 109, 490 individuals (9,824 were cases and 99, 666 were controls) from Vanderbilt University Medical Center’s (VUMC) de-identified EHRs. American College of Cardiology and the American Heart Association (ACC/AHA) Pooled Cohort Risk Equations had areas under receiver operating characteristic curves (AUROC) of 0.732 and areas under receiver under precision and recall curves (AUPRC) of 0.187. LSTM, CNN and GBT with temporal features achieved best results, which had AUROC of 0.789, 0.790, and 0.791, and AUPRC of 0.282, 0.280 and 0.285, respectively. The late fusion approach achieved a significant improvement for the prediction performance.

**Conclusions:** Machine learning and deep learning with longitudinal features improved the 10-year CVD risk prediction. Incorporating genetic features further enhanced 10-year CVD prediction performance, underscoring the importance of integrating relevant genetic data whenever available in the context of routine care.

## INTRODUCTION

Cardiovascular disease (CVD) is the leading cause of morbidity and mortality, accounting for one-third of all global deaths [1,2]. There have been several proposed several prediction models, including the Framingham risk score [3], American College of Cardiology/American Heart Association (ACC/AHA) Pooled Cohort Risk Equations [4], and QRISK2 [5]. These models are typically built upon a combination of readily-available cross-sectional risk factors such as hypertension, diabetes, cholesterol, age, and smoking status. Although the importance of conventional models cannot be ignored, well-known clinical risk factors for CVD explain only 50-75% of the variance in major adverse cardiovascular events [6]. About 15%-20% of patients who experienced myocardial infarctions had only one or two of these traditional risk factors and were not identified as being at “risk” of CVD according to current prediction models [7]. Given the fact that CVD is preventable, and that its first manifestation may be fatal, a new strategy to enhance risk prediction beyond conventional factors is critical for public health.

Electronic health records (EHRs) contain a wealth of detailed clinical information and provide several distinct advantages for clinical research, including cost efficiency, a large amount of data, and the ability to analyze data over time. Since its wide implementation in the United States, accumulated EHR data has become an important resource for clinical studies. [8]. Meanwhile, the recent convergence of two rapidly developing technologies—high-throughput genotyping and deep phenotyping within EHRs – presents an unprecedented opportunity to utilize routine healthcare data and genetic information to accelerate the improvement of healthcare. Many institutions and health care systems have been building EHR-linked DNA biobanks to enable such a vision. For example, Vanderbilt University Medical Center (VUMC), as of May 2018, has genotype data of over 50,000 individuals available for research.

Machine learning and deep learning approaches are particularly suited to the integration of big data, such as the data available within EHRs, especially when the EHR contains genetic information [9,10]. A recent study from the United Kingdom (UK) applied machine learning on conventional CVD risk factors from a large UK population and improved the overall prediction performance by 4.9% [11]. In the current study, we examined: i) the performance of machine learning and deep learning on longitudinal EHR data for the prediction of 10-year CVD risk, and ii) the benefits of incorporating extra genetic information.

## METHODS

### Study setting

We conducted the study using data derived from Synthetic Derivative, a de-identified copy of whole EHRs at VUMC. Synthetic Derivative maintains rich and longitudinal EHR data from over 3 million unique individuals, including demographic details, physical measurements, history of diagnosis, prescription drugs, and laboratory test results. As of May 2018, over 50,000 of these individuals have genotype data available.

We focused our analysis on individuals with European or African ancestry. We required individuals to meet the definitions of medical home [12]. We set the baseline date as 01/01/2007 to allow all individuals within the cohort to be followed-up for 10 years. For each individual, we split the EHR into: i) the observation window (01/01/2000 to 12/31/2006; 7 years) and, ii) the prediction window (01/01/2007 to 12/31/2016; 10 years). We extracted EHR data within the 7-year observation window to train a predictive model to predict CVD event occurred in prediction window.

Cases were individuals with ≥ 1 CVD diagnosis codes (the International Classification of Diseases, Ninth Revision, Clinical Modification [ICD-9-CM]: 411. * and 433. *) recorded within the 10-year prediction window. Controls were individuals without any ICD-9-CM code 411. * or 433. * during the 10-year prediction window.

### Study cohort

The study cohort included patients between the ages of 18 to 78 on 01/01/2000 (beginning of the observation window). Individuals with any CVD diagnosis (ICD-9-CM 411. * or 433. *) prior to the baseline date for the prediction window (i.e. 01/01/2007) were excluded. To reduce chart fragmentation and optimize the density of our longitudinal EHR data, we required that each individual to have at least one visit and at least one record of blood pressure measurement during the observation window [13,14]. We excluded inpatient physical or laboratory measures for all individuals.

In total, we identified 109, 490 individuals (9,824 cases and 99, 666 controls, mean [SD] age 47.4 [14.7] years; 64.5% female and 86.3% European) as our main study cohort. The case/control ratio was consistent with a previous report from a large EHR cohort [11]. Among these 109, 490 individuals, a subset of 10,162 individuals (2,452 cases and 7,710 controls) had genotype data available.

### Data preprocessing and feature extraction

Phenotypic data: we extracted features including demographics, variables used in the ACC/AHA Pooled Cohort Risk Equations (ACC/AHA Equations) (e.g. blood pressure measurements), physical measurements including BMI, and laboratory measures including glucose, triglyceride levels, and creatinine level (as a marker of renal function); such laboratory features have previously been reported relevant to CVD [11]. In addition, we applied chi-square (chi2) [15], a commonly used feature selection methods that can select independent features on EHR data and identified an additional 40 relevant diagnostic codes and medication codes (Table 1). Values for all features were extracted within the observation window.

**Table 1.**
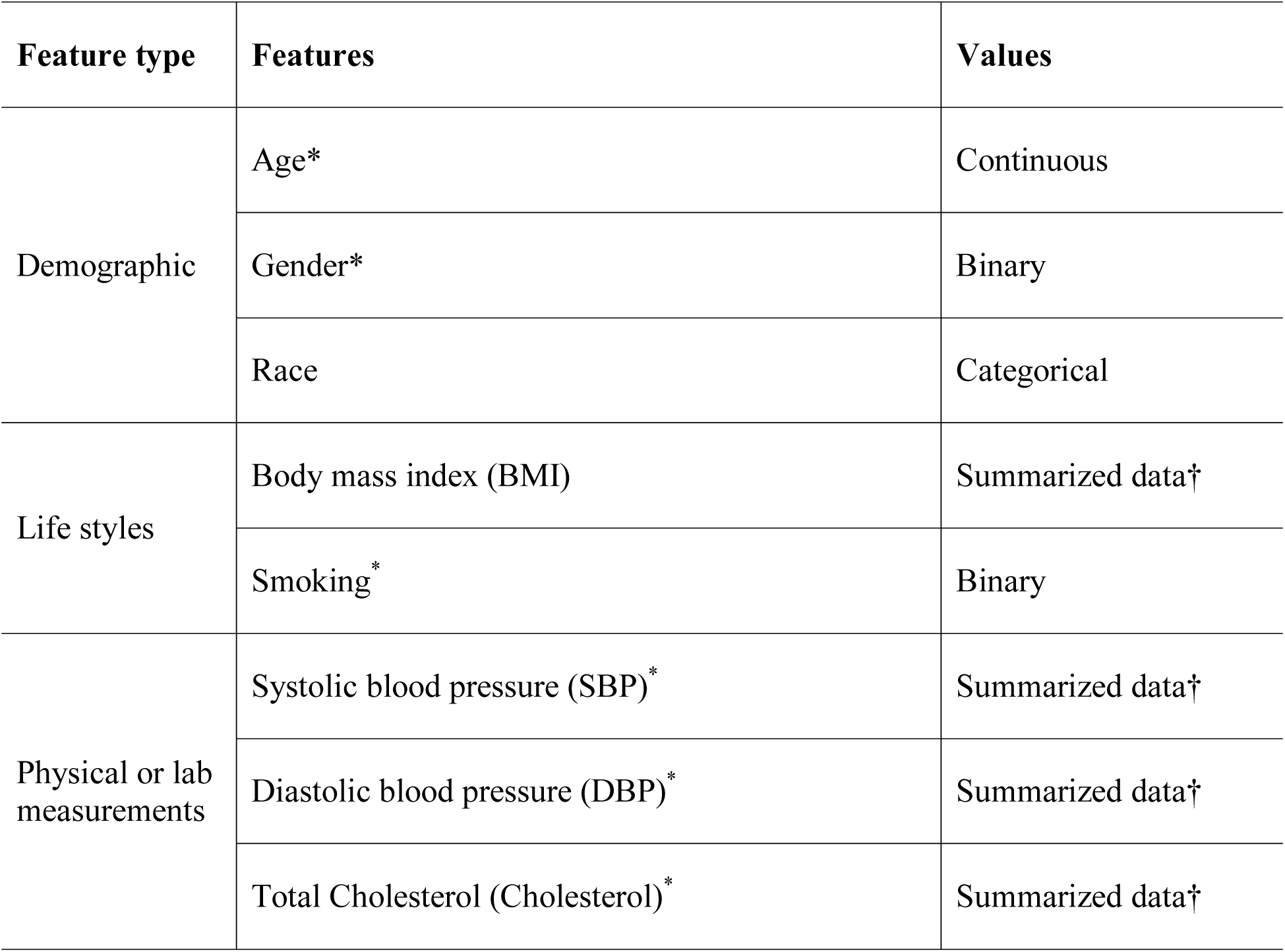
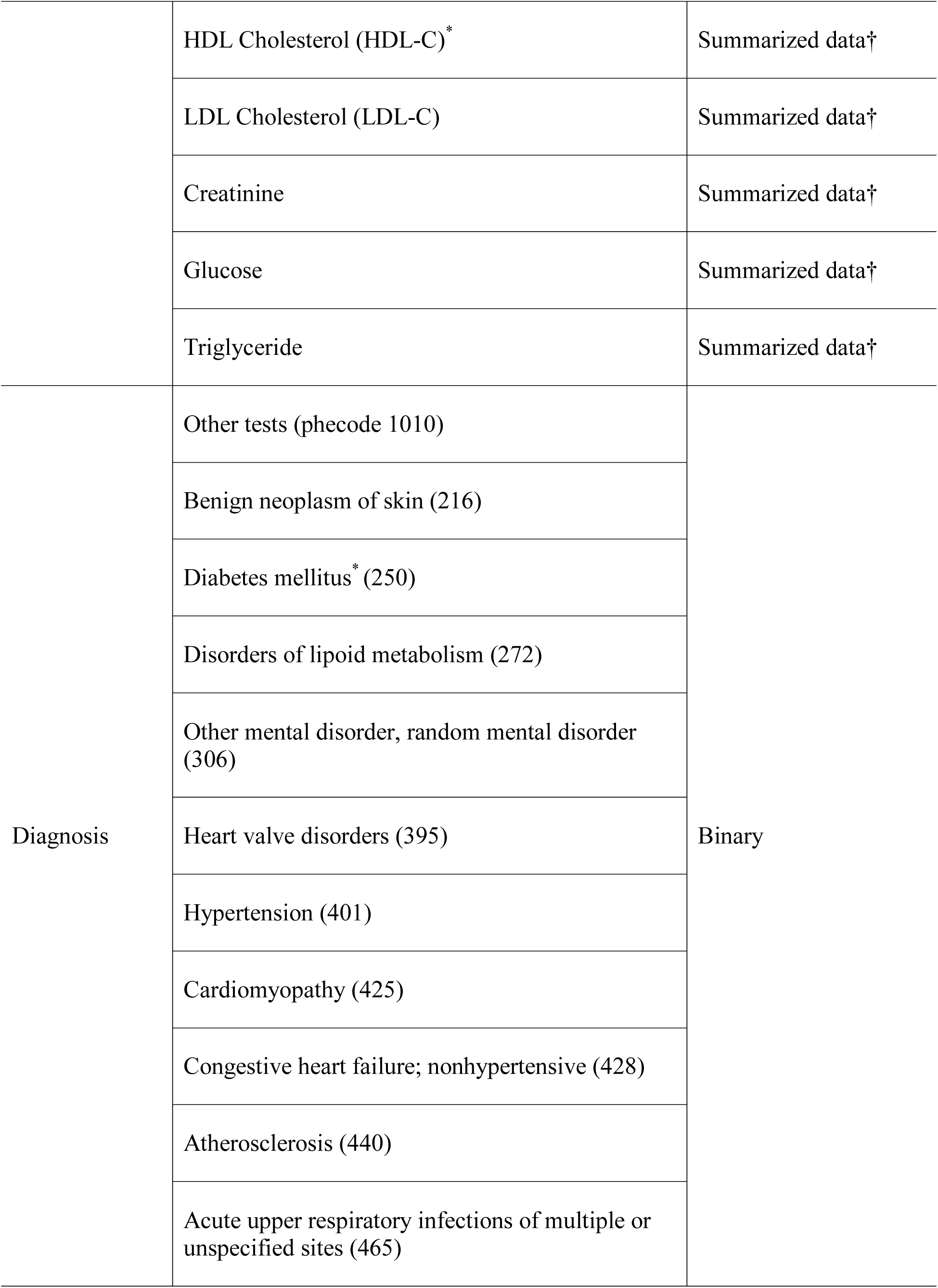
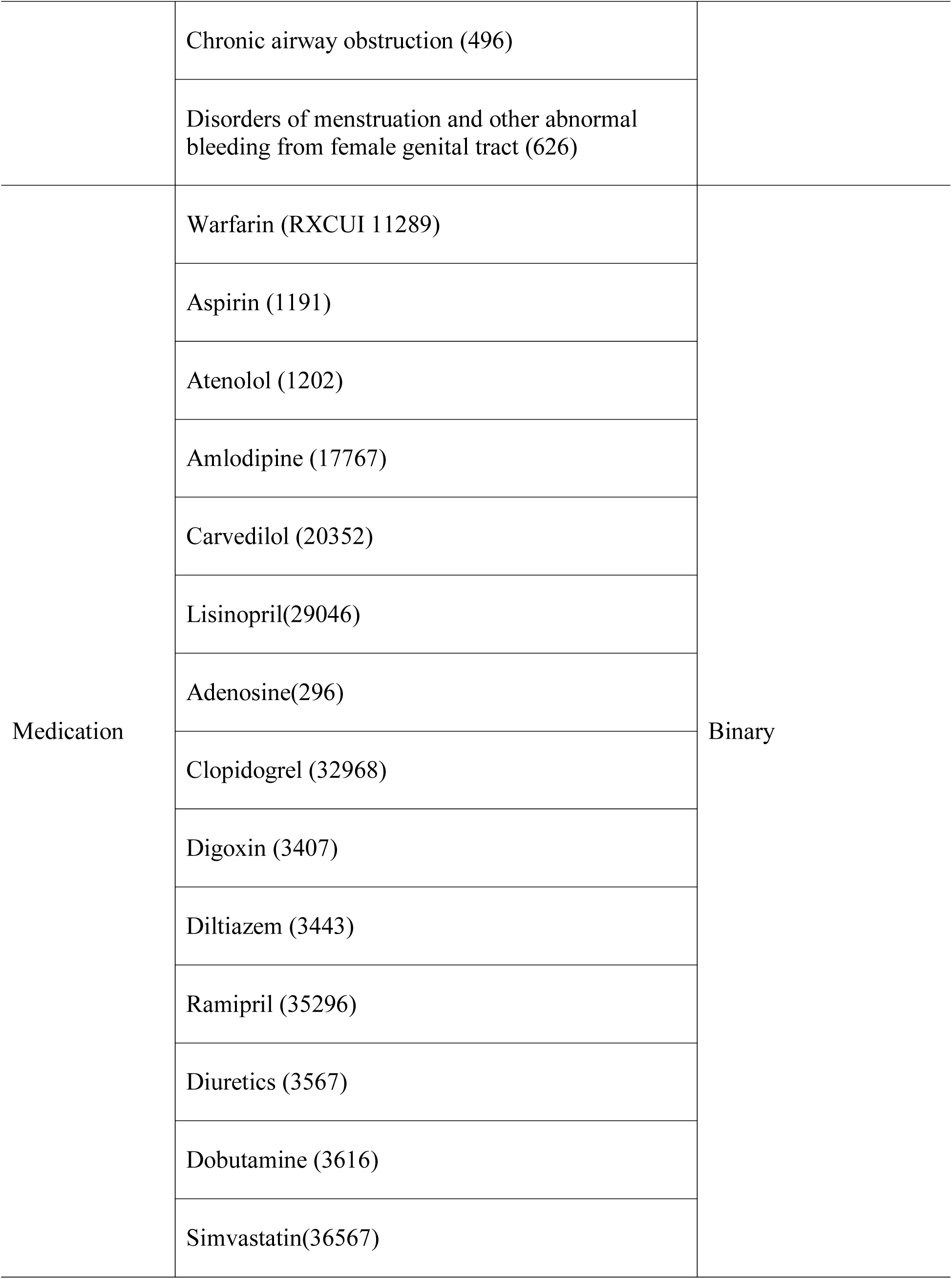
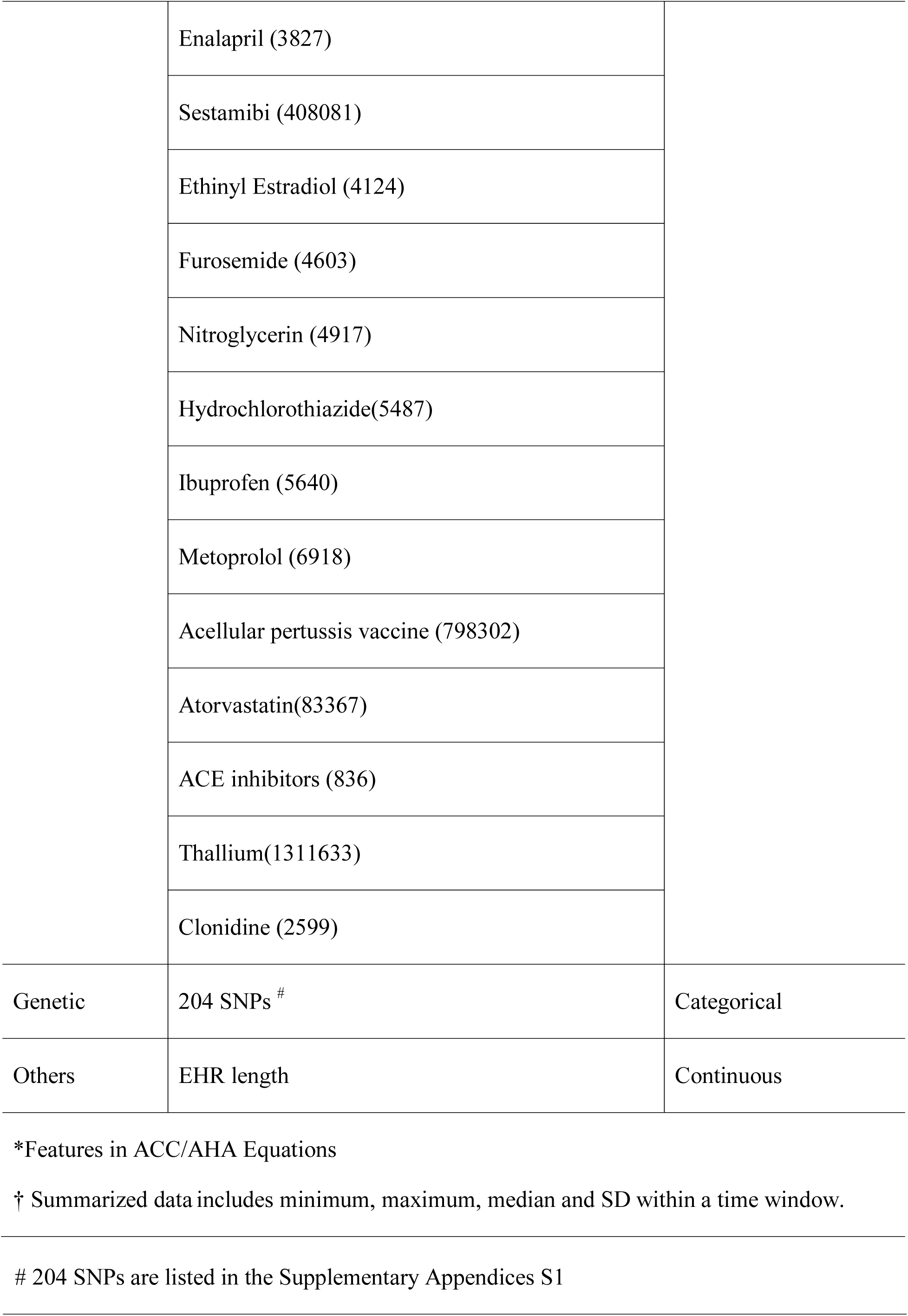
Features included in the machine-learning models.

We represented a physical measurement or laboratory feature with summarized data, e.g. minimum, maximum, median, and standard deviation (SD). We removed the outliers (>5 SD from the mean) to avoid unintended incorrect measurements (e.g. using lb. instead of kg. for body weight) [16]. If an individual had no such measure available within the EHR, we imputed the missing value with the median value of the group with the same age and gender [17]. We also added a dummy variable for each measure to indicate whether the test value was imputed.

For disease phenotypes, we followed a standard approach and grouped relevant ICD codes into distinct phecodes [19]. For medications, we collapsed brand names and generic names into groups by their composition (ingredients) and represented the groups using the RxNorm [19] concepts (RxCUIs) for this variable. For example, ‘Tylenol Caplet, 325 mg oral tablet’ and ‘Tylenol Caplet, 500 mg oral tablet’ were both mapped to ‘Acetaminophen’ (RxCUI 161). We used a binary value to indicate whether or not an individual had each diagnosis or prescription.

For genetic data, we selected 248 single nucleotide polymorphisms (SNPs) that have been previously reported to be associated with CVD in two large meta-analyses [20,21]. Among these SNPs, genotype data were available for 204 SNPs in our cohort and were included as features. Each SNP had a value 0, 1, or 2 based on the count of minor alleles for an individual. Table 1 shows the features that we used in the machine learning models.

### Experiment

#### Gold standard

We chose ACC/AHA Pooled Cohort Risk Equations for 10-year CVD risk as our baseline. For physical measurements or laboratory features (i.e. SBP/DBP and high-density lipoprotein [HDL]-cholesterol level), we used the most recent values prior to the split date, 01/01/2007.

Machine learning and deep learning with longitudinal EHR data to predict 10-year CVD risk (Experiment I)

The objective of this experiment is to examine 1) predictive performance achieved by machine learning and deep learning with longitudinal EHR data compared to golds standard, and 2) two different approaches we use to model the longitudinal EHR data for machine learning models (Figure 1).

**Figure 1.**
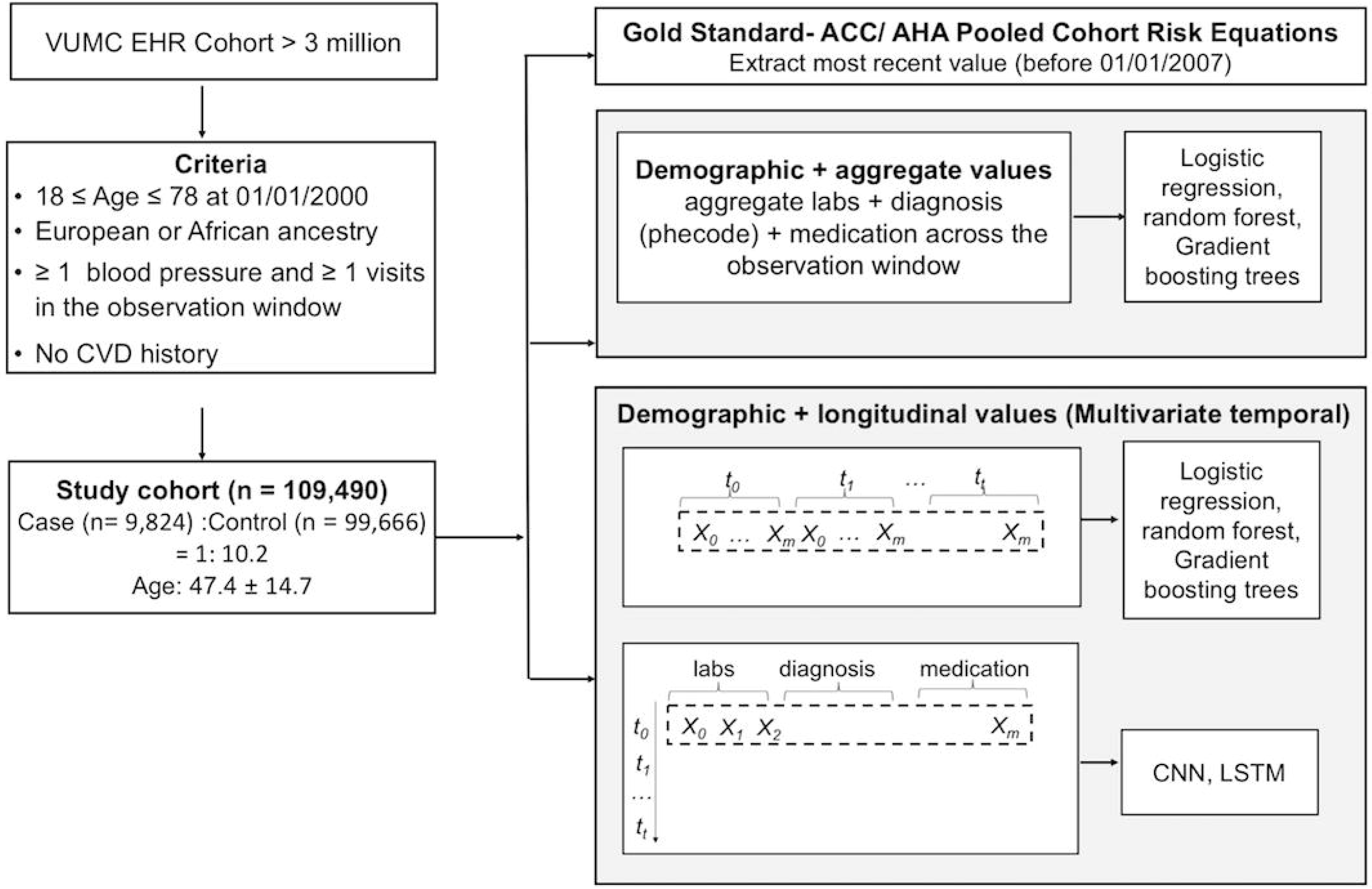
Flowchart of Experiment I: comparison of machine learning and deep learning models on longitudinal features against baselines.

#### Aggregate features

We aggregated features across the 7-year observation window (e.g. median, max, min and SD of HDL from 01/01/2000 to 12/31/2006).

#### Multivariate temporal features

We exploited the temporal information in the longitudinal EHR data by dividing the whole observation window into one-year slice window. Specifically, for physical or laboratory features, we extracted the median, max, min and SD values within one-year slice window. We replaced the missing physical or laboratory measures with the individual’s measurement on the closest date, e.g. using the HDL cholesterol result on 12/20/2005 instead if the individual had no HDL test in 2006. For diagnosis and medication features, we used a binary value to indicate whether or not an individual had each diagnosis or prescription in one-year slice window.

#### Machine learning and deep learning models

Three machine learning models, LR, RF and GBT were used in both aggregate and temporal features. Two deep learning models, Convolutional Neural Networks (CNN) [22] and Recurrent Neural Networks, using Long Short-Term Memory (LSTM) hidden units (LSTM) [23]) were applied to the temporal features. We compared their performance with the gold standard.

#### Implementation detail

We used CNN and LSTM on temporal features and concatenated an auxiliary input of demographic features to feed into a multilayer perceptron (MLP) with two hidden layers. More details can be found in Supplementary Appendices S2. LR, RF, and GBT models were implemented with Python Scikit-Learn 0.19.1 (http://scikit-learn.org/stable/) [24]. The CNN and LSTM models were implemented with Keras 2.1.3 (https://keras.io/) using Tensorflow1.6.1 as the backend.

#### Evaluation

We divided the dataset into a training and a test set with a 90/10 split and learned the models with a 10-fold stratified cross-validation using grid search on the training set. Finally, we evaluated the optimized model on the test set using area under a receiver operating characteristic curve (AUROC) and average precision, also known as area under precision-recall curve (AUPRC) [25]. For each machine learning model, we repeated the above processed 10 times. For deep learning models, we randomly divided the data into training, validation, and testing sets with a ratio of 8:1:1 and iterated the process for 10 times. We reported the mean and SD of AUROC and AUPRC.

Machine learning and deep learning with additional genetic information to predict 10-year CVD risk. (Experiment II)

The objective of this experiment is to examine combining genetic features with demographic and longitudinal EHR data compared to only using demographic and longitudinal EHR data for 10-year CVD prediction. To meet the objectives, we used a subset of 10,162 had genotyped data from the main study cohort of 109, 490 individuals. It is also a subset of BioVU (VUMC’s de-identified DNA biobank) that contains nearly >50,000 genotyped individuals.

We developed a two-stage framework of using late-fusion approach to incorporate EHR and genotyped features. Late-fusion is an effective approach to enhance prediction accuracy by combining the prediction results of multiple models trained separately by a group of features. [26] Here, we trained two machine learning models separately by EHR data and genotyped data and used a subset of 10,162 which had both available EHR and genotyped data to train and test a fusion model based on the prediction results. (Figure 2 and 3). The subset of 10,162 individuals (intersect cohort) was randomly split into a training set (8,129 individuals) and a holdout test set (2, 033 individuals) with an 80/20 split. The training set is used for training the fusion model at final decision level. The holdout test set is used for comparing the performance of models trained with only EHR data and the proposed late-fusion approach.

**Figure 2.**
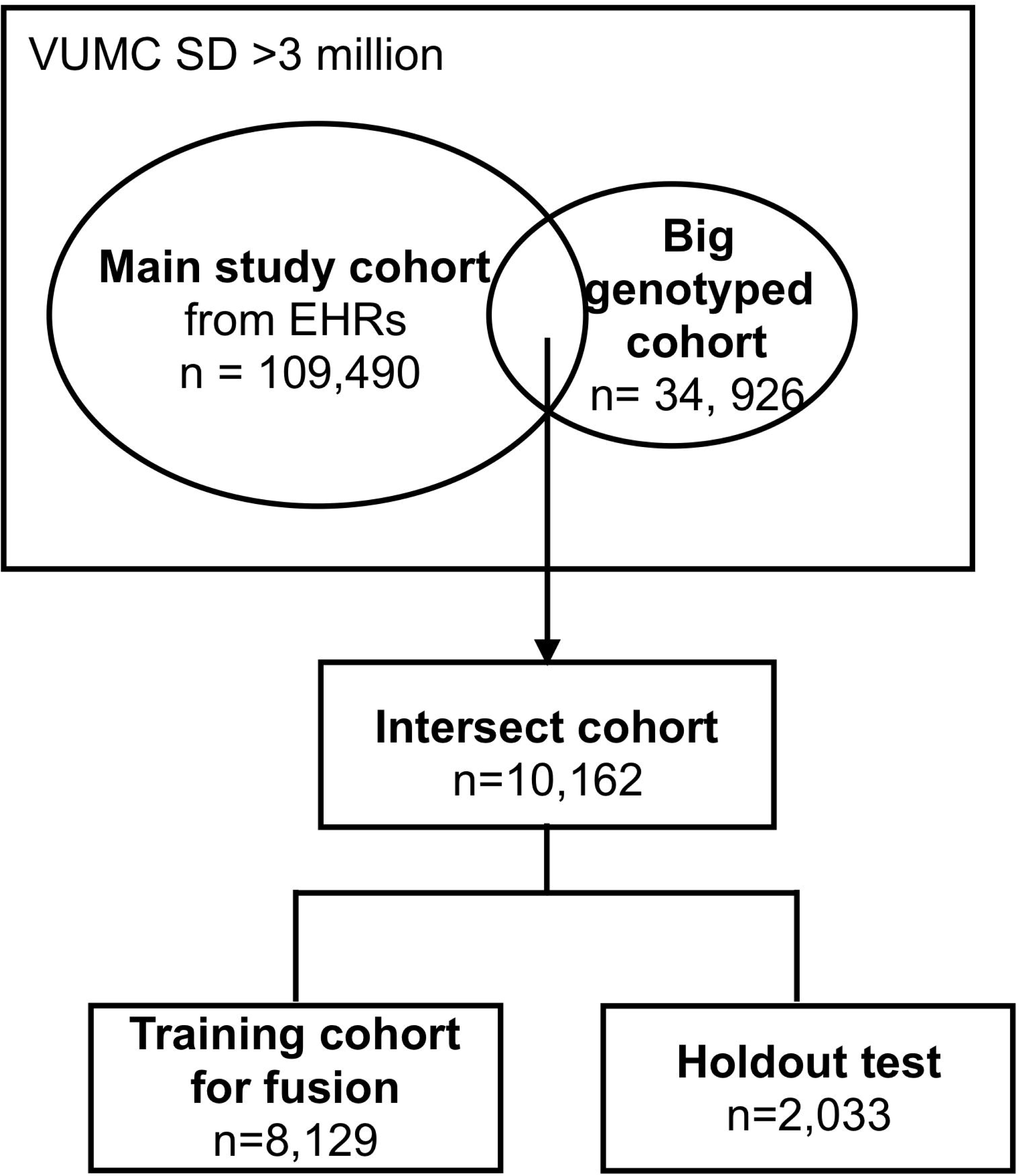
Flowchart of selecting cohort for late-fusion approach

**Figure 3.**
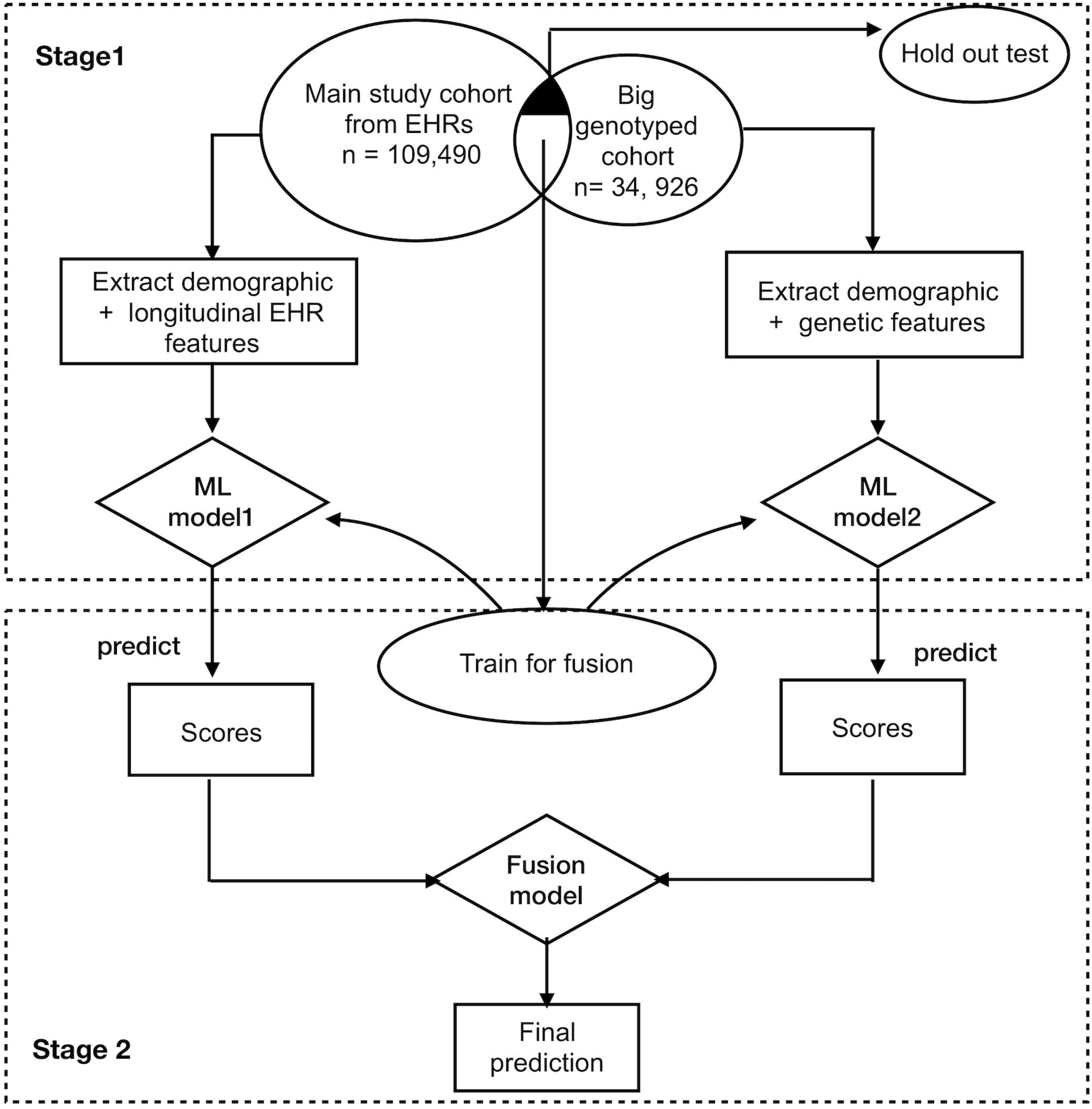
Framework for proposed late fusion approach to combine the genetic features with longitudinal EHR features.

In the first stage of the framework, we trained a machine learning model (model1) with longitudinal EHR features on the main study cohort (removing holdout test set). We trained another machine learning model (model 2) with 204 SNPs features on a big 34,926 genotyped cohort (removing holdout test set), which shared similar criteria with the main study cohort except for not restricting to the criteria for having >1 record of SBP in the observation window. In the second stage, we combined the predictions scores of two models on the training set (8, 129 individuals) to train a late fusion model. We used gradient boosting trees for the model 1 because it has good generalizability as an ensemble approach to make it more robust. We used the logistic regression as model 2 and the late fusion model.

To compare the performance of adding genetic features, we evaluated prediction performance of model 1 and fusion model on the holdout test set (2,033 individuals). We performed 5-fold cross-validation and repeated the process 10 times. We reported the mean and SD of AUROC and AUPRC.

## RESULTS

### Machine learning and deep learning models with longitudinal EHR data to predict 10-year CVD risk (Experiment I)

Table 2 shows the results for the experiment. The performance of the gold standard (AUROC 0.732, AUPRC 0.187) was consistent with other study reports [11,27]. Compared with gold standard, all three machine-learning models with aggregate features achieved significant improvements over the prediction metrics. For AUROC, RF increased the performance from 0.732 to 0.765, an absolute (relative) improvement of +0.033 (+4.5%). LR [+0.044 (+ 6.0%)] and GBT [+0.05 (+6.8%)] had a higher increase rate. For AUPRC, the improvement was much bigger, from RF [0.059 (+31.6%)] to GBT [+ 0.081 (+43.3%)].

**Table 2.** Performance of machine learning and deep learning models predicting 10-year CVD risk. The + indicates that the mean is significantly different from the mean of gold standard (p < 0.05), when evaluated using the *t*-test. # indicates that the mean of the model on longitudinal one-year slice window is significantly different from the model with aggregate features.

| Method | AUROC | AUPRC |
| --- | --- | --- |
| ACC/AHA Equations | 0.732 (± 0.010) | 0.187 (± 0.009) |
| **Machine learning models on aggregate features across seven-year window** |  |  |
| Logistic regression (LR) | 0.776 (± 0.008)<sup>+</sup> | 0.260 (± 0.014)<sup>+</sup> |
| Random forest (RF) | 0.765 (± 0.009)<sup>+</sup> | 0.246 (± 0.009)<sup>+</sup> |
| Gradient boosting trees (GBT) | 0.782 (± 0.009)<sup>+</sup> | 0.268 (± 0.014)<sup>+</sup> |
| **Machine learning models on longitudinal features within one-year window (temporal)** |  |  |
| Logistic regression (LR) | 0.781 (± 0.007)<sup>+</sup> | 0.273 (± 0.013)<sup>+#</sup> |
| Random forest (RF) | 0.753 (± 0.008)<sup>+</sup> | 0.236 (± 0.010)<sup>+</sup> |
| Gradient boosting trees (GBT) | 0.791 (± 0.008)<sup>+#</sup> | 0.285 (± 0.013)<sup>+#</sup> |
| **Deep learning models on longitudinal features within one-year window (temporal)** |  |  |
| LSTM | 0.789 (± 0.011)<sup>+</sup> | 0.282 (± 0.012)<sup>+</sup> |
| CNN | 0.790 (± 0.012)<sup>+</sup> | 0.280 (± 0.012)<sup>+</sup> |

Compared to the aggregate features, using longitudinal features further improved the prediction performance across most models. AUROC of GBT is improved from 0.782 to 0.791 [+ 0.009 (+1.2%)] and the AUPRC of GBT is improved from 0.268 to 0.285 [+0.017; (+6.3%)]. LR [+0.0013 (5.0%)] also had a significant improvement in AUPRC. For deep learning models with longitudinal features, LSTM and CNN achieved nearly same results as GBT, better than the LR and RF. Overall, the best result achieved by GBT using longitudinal features increased the AUROC of gold standard +0.059 (+8.1%) and AUPRC +0.098 (+52.4%).

#### Feature importance

We listed top features for each of optimized machine learning models in Table 3. Feature importance was determined by the coefficient effect size from the LR model. For RF and GBT, which are based on decision-trees, the features are ranked according to the impurity (information gain/entropy) decreasing from each feature. Since CNN and LSTM are black box models, estimation of each feature’s contribution to predict CVD risk is difficult, so we were not able to analyze the feature importance of the deep learning models in this study.

**Table 3.** Top 10 features for machine learning prediction in descending order of coefficient effect size or feature importance returned by RF and GBT. Systolic Blood Pressure (SBP); Diastolic Blood Pressure (DBP).

| **LR with aggregate features** | **RF with aggregate features** | **GBT with aggregate features** | **LR with longitudinal features** | **RF with longitudinal features** | **GBT with longitudinal features** |
| --- | --- | --- | --- | --- | --- |
| EHR length | EHR length | Age | EHR length | EHR length | Age |
| Max LDL-C | Age | EHR length | Age | Age | EHR length |
| Min Creatinine | Max BMI | SD Creatinine | SD Glucose in 2000 | Aspirin in 2006 | Smoking |
| Age | Min BMI | Smoking | SD Creatinine in 2000 | Max SBP in 2006 | Heart valve disorders in 2006 |
| Max HDL-C | Median BMI | Min BMI | Max HDL-C 2005 | Min BMI in 2006 | Hypertension in 2006 |
| Max BMI | Max SBP | Heart valve disorders (Phecode 395) | SD Glucose in 2006 | Median BMI in 2005 | Aspirin in 2006 |
| Max Cholesterol | Median SBP | Min Glucose | Median LDL-C in 2006 | Median SBP in 2006 | Disorders of lipid metabolism in 2006 |
| Max DBP | SD BMI | Max SBP | Median BMI in 2006 | Max BMI in 2006 | Clopidogrel in 2006 |
| Median Trigs | MIN SBP | Max Trigs | Median Cholesterol in 2006 | Min BMI in 2001 | Max SBP in 2006 |
| Min Cholesterol | Max DBP | Aspirin | Heart valve disorders in 2006 | Min BMI in 2002 | SD Glucose in 2006 |

The conventional risk factors such as age, blood pressure and total cholesterol were consistently present as top 10 features in all three machine learning models. BMI, Creatinine and Glucose that were not in ACC/AHA equations were determined as important features in machine learning models. Moreover, the maximum, minimum, and SD of laboratory values showed promising contributions to the models. GBT models preferred diagnoses such as heart valve disorder, hypertension, and lipid disorders over other features.

For machine learning models with longitudinal features, LR models selected laboratory values in the years 2000 and 2006 (e.g. SD Glucose in 2000 and 2006). The RF models chose BMI in multiple years. Whereas GBT models prioritized the medical conditions in the most recent year (year 2006) in the observation window.

### Evaluate incorporating genetic features for machine learning models to predict 10-year CVD risk (Experiment II)

Table 4 reported the results of Experiment II. GBT with only longitudinal EHR features improved AUROC of gold standard from 0.698 to 0.710 [+0.012 (+1.7%)] and AUPRC from 0.396 to 0.427 [+0.031(+7.8%)]. The proposed late fusion approach for adding genetic features further improved the metrics, with AUROC +0.015 (+2.1%) and AUPRC +0.036 (+9.1%).

**Table 4.**
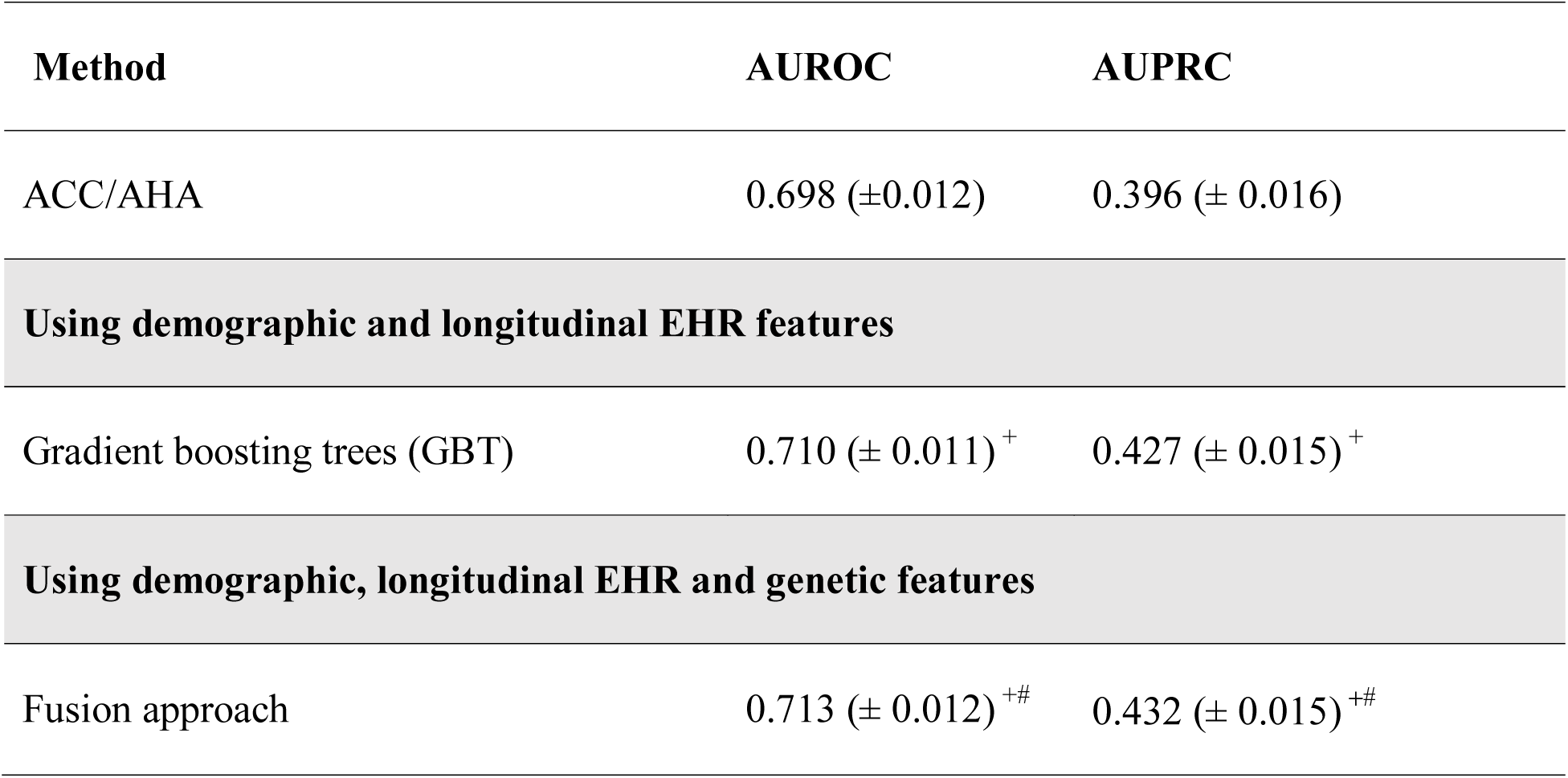
Comparison of predicting 10-year CVD risk with genetic features and without genetic features. + indicates that the mean is significantly (p < 0.05) different from gold standard, and # indicates that the mean is significantly different from GBT using demographic and longitudinal EHR features, when evaluated using the paired *t*-test.

We listed the top ten features in the pre-trained model with genetic data in Supplementary Appendices S3. SNP (rs2789422) was ranked as the second most important feature after age.

## DISCUSSION

Our results demonstrate that machine learning models with longitudinal EHR information can improve the prediction of 10-year CVD risk. We also showed that incorporating genetic data can enhance 10-year CVD risk prediction.

We used a large dataset including longitudinal EHR information of 109, 490 individuals. The prediction result of ACC/AHA (AUROC of 0.732, AUPRC of 0.187) was consistent with previous studies (AUROC of 0.728 in a study conducted in the UK) [11].

For machine learning models with aggregate values, as we used summarized data for physical and laboratory features, and we also included 40 additional pre-selected features including diagnosis codes and medication codes, the performance of prediction was significantly improved. Further, the min, max and SD values were ranked higher in importance than the median values. BMI, medications (e.g. aspirin) that were not used by the ACC/AHA equations were also present in the top 10 features.

Longitudinal information reflects the fluctuation of physiological factors over time, which can be used for prediction models to enhance CVD risk prediction. The most recent results from the STABILITY trial suggested the higher visit-to-visit variabilities of both systolic and diastolic blood pressures are strong predictors of increased risk of CVD, independently of mean blood pressure[28]. By zooming in the observation window of one-year slice time, we constructed multivariate temporal features for machine learning models and deep learning models. The results showed that it improved the prediction performance. CNN and LSTM that allows for exhibiting dynamic temporal changes, outperformed LR and RF models. Surprisingly, GBT almost had similar performance as LSTM and CNN. The time steps (7 years, 7-time steps) may not be long enough to activate the gates of LSTM. Another reason is that a 10-year follow-up prediction window may be a little long thereby removing the advantage of LSTM and CNN in capturing the dependency with the observation and prediction.

Our approach also underscores the importance of including genetic variants. It has long been known that CVD has a sizeable hereditary component [3], and emerging data continue to increase our understanding of the genetic architecture underlying this important clinical trait [20,21]. Previous studies have uncovered many novel genetic associations with CVD for risk factors that are also heritable such as lipids, blood pressure, and diabetes [29,30]. While polygenic scores have been used to summarize genetic effects for diseases, strategies to combine genetic variants with other biological and lifestyle factors for existing predictive models remains a topic of intense ongoing investigation. Although 10,162 individuals (2,452 cases and 7,710 controls) of our main study cohort had genotype data available, this subset may still limit our power for large scale genetic analyses and machine learning. Since the quality of prediction often depends on the amount of available training data, without sufficient training data, the learning models cannot differentiate useful patterns from noise and predictive accuracy may underperform. To overcome this challenge, we proposed a late-fusion approach to pre-train the models with EHR features and genetic features separately by taking advantage of a larger genotyped cohort (34,926).

From the results, we can see that adding genetic features offered benefit to clinical features and significantly improved the performance compared to gold standard and only using longitudinal EHR features.

### Importance of Genetic Features

We present the top 10 features identified from the cohort (Supplementary Appendices S3). Age remains the strongest predictor for CVD (coefficient 0.747), followed by gender, EHR length and two variants from *MIA3* gene. Although dyslipidemia is one of the most important risk factors for CVD, none of the top genes was strong predictor for circulating lipid levels, except *LPA* gene.

While *LPA* genotype are associated with circulating lipid levels, it also strongly influenced Lp(a) levels which was an independent CVD predictor with or without statin treatment [31]. For decades, lipid-lowering medications (especially statins) have been shown to be effective in both primary and secondary CVD prevention. Our observations highlight the importance of CVD risk determinants independent of lipid levels. These findings underscore the importance of targeting residual CVD risk through non-lipid mechanisms.

We acknowledge the limitations that, 1) we manually abstracted a subset of the physical or laboratory features known to impact CVD risk, and we planned to incorporate more laboratory features that could be automatically selected by feature engineering from the EHR, and 2) we only used 204 SNPs in our study, whereas some of effects of the SNPs are modeled by phenotypes (e.g., a SNPs affecting cholesterol are better captured by direct cholesterol measurements). Yet some SNPs for *endophenotypes* are more predictive of CVD events than the endophenoytpe itself [31]. As each SNP has a relatively small effect size compared with other features like age, gender, and diabetes, and thus may not contribute much to the predictive ability of the models, we believe that with more phenotypic and genetic information available in larger cohorts may further improve prediction. This study confirmed that combining phenotypic and genetic information with robust computational models can improve disease prediction.

## FUNDING

The project was supported NIH grant P50 GM115305, R01 HL133786, R01 GM120523, T32 GM007347 from the National Institute of General Medical Studies for the Vanderbilt Medical-Scientist Training Program, and T15 LM007450 from the National Library of Medicine for the Vanderbilt Biomedical Informatics Training Program.

The dataset used in the analyses described were obtained from Vanderbilt University Medical Centers BioVU which is supported by institutional funding and by the CTSA grant ULTR000445 from NCATS/NIH. Genome-wide genotyping was funded by NIH grants RC2GM092618 from NIGMS/OD and U01HG004603 from NHGRI/NIGMS."

The dataset(s) used for the analyses described were obtained from Vanderbilt University Medical Center’s BioVU which is supported by institutional funding, the 1S10RR025141-01 instrumentation award, and by the CTSA grant UL1TR000445 from NCATS/NIH. The authors wish to acknowledge the expert technical support of the VANTAGE and VANGARD core facilities, supported in part by the Vanderbilt-Ingram Cancer Center (P30 CA068485) and Vanderbilt Vision Center (P30 EY08126).

This study was supported by GM120523, GM109145, HL133786, 5T32GM080178-09, K23AR064768, Rheumatology Research Foundation (K-supplement), American Heart Association (16SDG27490014 and 15MCPRP25620006), HG008672, R01 LM010685, GM115305 and Vanderbilt Faculty Research Scholar Fund. The dataset used for the analyses described were obtained from Vanderbilt University Medical Center’s resources, BioVU and the Synthetic Derivative, which are supported by institutional funding and by the Vanderbilt National Center for Advancing Translational Science grant 2UL1 TR000445-06 from NCATS/NIH. Existing genotypes in BioVU were funded by NIH grants RC2GM092618 from NIGMS/OD and U01HG004603 from NHGRI/NIGMS. The funders had no role in study design, data collection and analysis, decision to publish, or preparation of the manuscript.

## COMPETING INTEREST

The authors have no competing interests to declare.

